# The cytosolic cysteinyl-tRNA synthetase is capable of regulating ATF4 translation independent of eIF2α phosphorylation state

**DOI:** 10.1101/2024.12.23.630120

**Authors:** Acer E. Xu, Jonathan S. Weissman

## Abstract

In mammalian cells, the integrated stress response (ISR) triggers ATF4 translation under general conditions of cell stress. Mammalian target of rapamycin (mTOR) signaling also triggers ATF4 translation under general pro-growth conditions. While the transcriptomic changes of these two contradictory pathways are have been studied, a full understanding of all pathways capable of increasing ATF4 translation and how these pathways are regulated and interact with each other remains unknown. In a genome-wide CRISPRi screen, we found that loss of CARS is sufficient to activate ATF4 translation independent of canonical ISR signaling. This ATF4 translation does not require eIF2α phosphorylation and does not require MTOR kinase activity. In a genome-wide epistasis CRISPRi screen using a staggered sgRNA infection strategy, we identified METAP2 and OGT as potential downstream factors in this pathway. This represents a novel pathway for activation of ATF4 translation that may enable targeted manipulation of ATF4 for beneficial therapeutic outcomes.

## Introduction

The maintenance of cellular homeostasis requires cells to be able to respond to changes to their environment (Hartl, 2016). Cells must be able to take advantage of situations favorable to growth, and respond to stresses that cause disruption to essential functions by changing gene expression to restore balance. The integrated stress response (ISR) is an evolutionarily conserved intracellular signaling network that allows cells, tissues, and organisms to adapt to environmental changes (Pakos-Zebrucka *et al*., 2016; Pavitt, 2018; Costa-Mattioli and Walter, 2020).

The ISR regulates homeostasis by reprogramming gene expression. Four specialized kinases (PERK, GCN2, PKR and HRI) sense various stresses that result in the phosphorylation of a single serine on translation initiation factor eIF2α (eukaryotic translation initiation factor 2 subunit alpha, gene name *EIF2S1*) (Levin *et al*., 1980; Dever *et al*., 1992; Harding *et al*., 1999; Han *et al*., 2001; Hinnebusch, 2005; Costa-Mattioli and Walter, 2020; Wu *et al*., 2020). This phosphorylation of eIF2α inhibits the activity of the eIF2α guanine nucleotide exchange factor eIF2B, leading to a decrease in protein translation initiation rates (Kashiwagi *et al*., 2019; Kenner *et al*., 2019). However, this phosphorylation of eIF2α also activates the translation of specific mRNAs, including the key transcription factor activating transcription factor 4 (ATF4). The ATF4 mRNA contains short inhibitory upstream open reading frames (uORF) in its 5′-untranslated region that prevents translation initiation at the canonical AUG (Vattem and Wek, 2004). When eIF2α is phosphorylated, these uORFs are bypassed and ATF4 is translated (Lu *et al*., 2004). By upregulating the synthesis of specific transcription factors and downregulating global protein translation, the ISR aims to restore homeostasis. If the stress cannot be mitigated, prolonged ISR signaling triggers apoptosis and cell death (Zinszner *et al*., 1998; Costa-Mattioli and Walter, 2020).

It is also clear that the ISR is implicated in the pathology of several disorders, with regulation of the ISR shown to be a potential therapeutic avenue for the treatment of a variety of diseases (Lu *et al*., 2004; Boyce *et al*., 2005; Rosen *et al*., 2009; Bryk *et al*., 2011; Sidrauski *et al*., 2013, 2015). For example, genetic and pharmacological evidence has shown that tuning the ISR improves recovery in mouse models of concussions (Chou *et al*., 2017). Similarly, the ISR has been implicated in many other complex diseases, including cancer and metabolic disorders, with both reduced and increased ISR activation showing maladaptive phenotypes depending on cellular and disease contexts (Costa-Mattioli and Walter, 2020). Therefore, ISR activation or inhibition to restore homeostasis and optimal cell fitness will depend on the disease and phenotype.

Recent studies have shown that beyond its role as the major downstream effector of the ISR, ATF4 translation can also be activated by pro-growth signals that lead to mammalian target of rapamycin (mTOR) signaling, a growth signaling pathway regulated by MTOR (mechanistic target of rapamycin kinase, gene name *MTOR*) (Wang and Proud, 2008; Park *et al*., 2017; Torrence *et al*., 2021). Despite these two pathways being activated by seemingly contradictory environmental cues, both pathways converge on ATF4 activation, with MTOR-mediated activation of ATF4 occurring independently of the ISR and phosphorylation of eIF2α (Park *et al*., 2017). However, the MTOR-ATF4 dependent gene program represents only a small and partially overlapping subset of the genes upregulated by the ISR-ATF4 program (Torrence *et al*., 2021).

The full accounting of pathways capable of activating ATF4 translation, as well as how different ATF4 activating pathways result in distinct gene expression signatures, remains unknown. Here, we report that inhibition of the cytosolic cysteinyl-tRNA synthetase (CARS, gene name *CARS1*) is capable of activating ATF4 translation in a manner independent of eIF2α phosphorylation. We find that this ATF4 translation results in a true transcriptional response similar to both canonical ISR and mTOR mediated ATF4 transcriptomes. Finally, we show that this *CARS1* knockdown mediated ATF4 activation may be dependent on the activity of METAP2 and OGT.

## Results

### A system for studying eIF2α phosphorylation-independent ATF4 translation

To monitor ATF4 activation, we introduced a previously published reporter for ATF4 translation (Guo *et al*., 2020) into K562 cells containing the machinery necessary for CRISPR interference. The reporter (thereafter referred to as the “ATF4-mApple reporter) was activated by various drug induced stress in K562 cells (Figure 1a, 1b) (Oslowski and Urano, 2011). We tested the role of the four known eIF2α kinases, and found that the ATF4-mApple reporter retained pathway specificity when exposed to stress, with PERK being required for ATF4-mApple reporter activation upon treatment with thapsigargin, a known inducer of endoplasmic reticulum stress in K562 cells that requires PERK kinase activity to induce ATF4 translation (Vattem and Wek, 2004; Replogle *et al*., 2022b) (Figure 1c).

**Figure 1:**
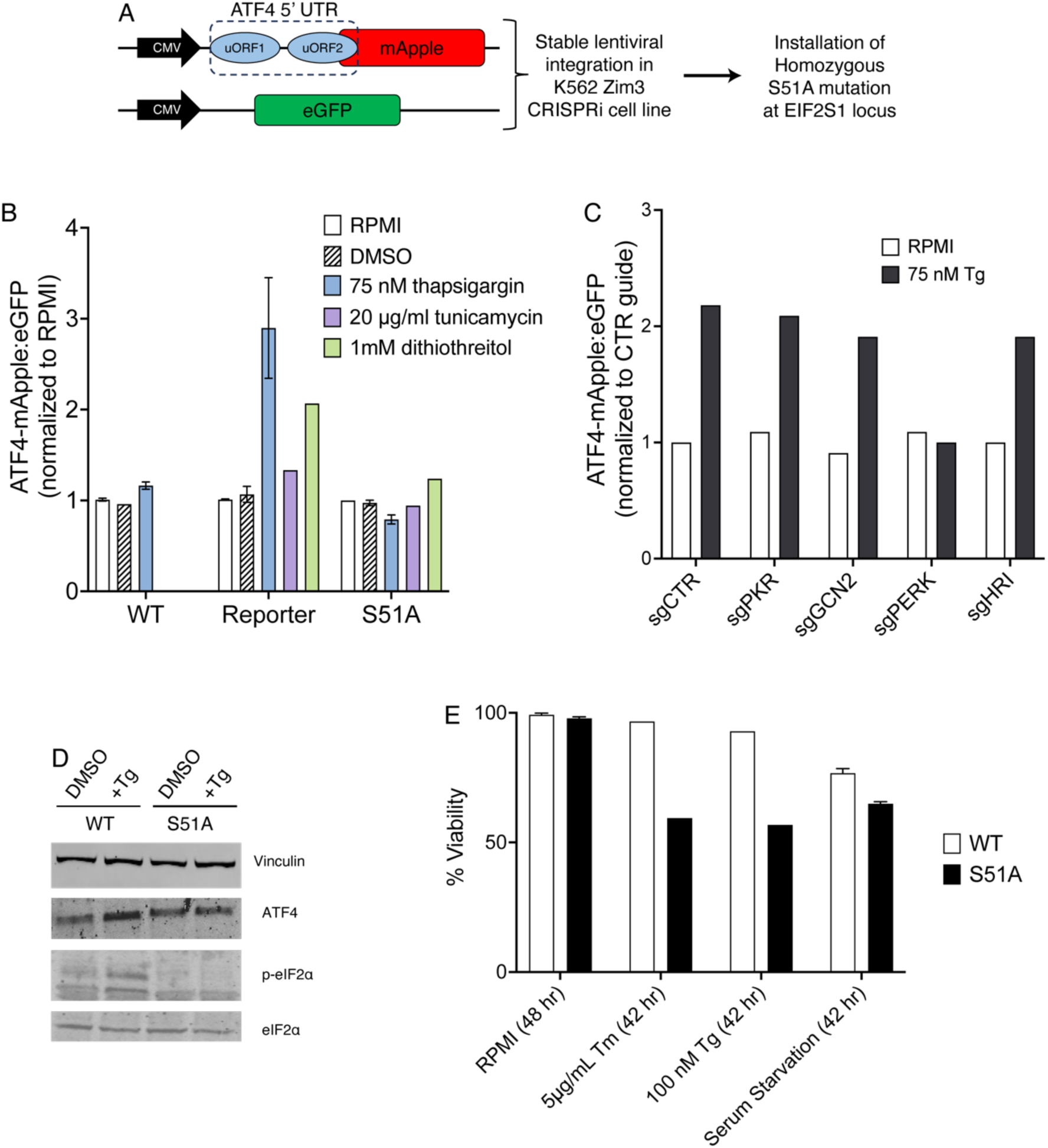
A system for studying eIF2α phosphorylation-independent ATF4 translation (A) Scheme for construction of ATF4 translation reporter, followed by introduction of homozygous S51A mutation. (B) Cells were exposed to 75 nM thapsigargin or vehicle (DMSO) for 16 hours, before measuring reporter levels by flow cytometry. (C) WT and S51A reporter cells expressing single sgRNAs were exposed to 75 nM thapsigargin or vehicle (DMSO) for 16 hours, before measuring reporter levels by flow cytometry. (D) Immunoblot of ATF4, eIF2α, and p-eIF2α. Cells were exposed to 75 nM thapsigargin or vehicle (DMSO) where indicated. (E) WT and S51A reporter cells were exposed to indicated drug or cell culture media containing no FBS for 42 hours, before measuring percent cells viable via flow cytometry.

It is well known that canonical activation of ATF4 translation via the four known eIF2α kinases requires the phosphorylation of the specific serine 51 residue on eIF2α (Pathak *et al*., 1988; Dever *et al*., 1992; Scheuner *et al*., 2001). To study if there are non-eIF2α phosphorylation dependent pathways for activating ATF4 translation, we used CRISPR cutting and subsequent homology directed repair to introduce a homozygous mutation in the genomic *EIF2S1* locus, changing the serine residue at position 51 to an alanine residue (Figure S1a). Homozygous S51A cells still express normal levels of eIF2α as measured by western blotting, but do not phosphorylate eIF2α when treated with thapsigargin, and do not upregulate ATF4 translation when exposed to protein folding stress (Figure 1d, S1b).

Previous studies using a mouse derived S51A fibroblast model have shown that S51A cells exhibit increased sensitivity to protein folding stress but not to general growth inhibition (Scheuner *et al*., 2001). In agreement with this, S51A cells are viable in standard cell culture conditions, but show increased sensitivity to chemically induced protein folding stress known to canonically induce the ISR (Figure 1e). Because ATF4 translation signaled through the mTOR pathway occurs independent of eIF2α phosphorylation, we hypothesized that S51A cells should still be responsive to growth cues signaled via this pathway. In line with this reasoning, we found that serum starvation of S51A and wild-type cells both led to similar levels of growth inhibition (Figure 1e).

### CRISPRi screen identifies CARS as an effector of ATF4 translation

To identify novel pathways capable of regulating ATF4 translation independent of eIF2α phosphorylation, we performed a genome-wide CRISPRi screen, sorting our screened cells by ATF4-mApple:eGFP ratio to identify genes that affected translational regulation of ATF4 when knocked down (Figure 2a, S2a). Generally, we found that the most represented class of genes that saw significant enrichment or depletion were components of translation initiation machinery (Figure S2b), a result in line with decades of biochemical work studying translation initiation (Lomakin and Steitz, 2013). As expected, strong hits that activated the ISR in this category were the components of the eIF2α and eIF2B complexes (Figure 2b).

**Figure 2:**
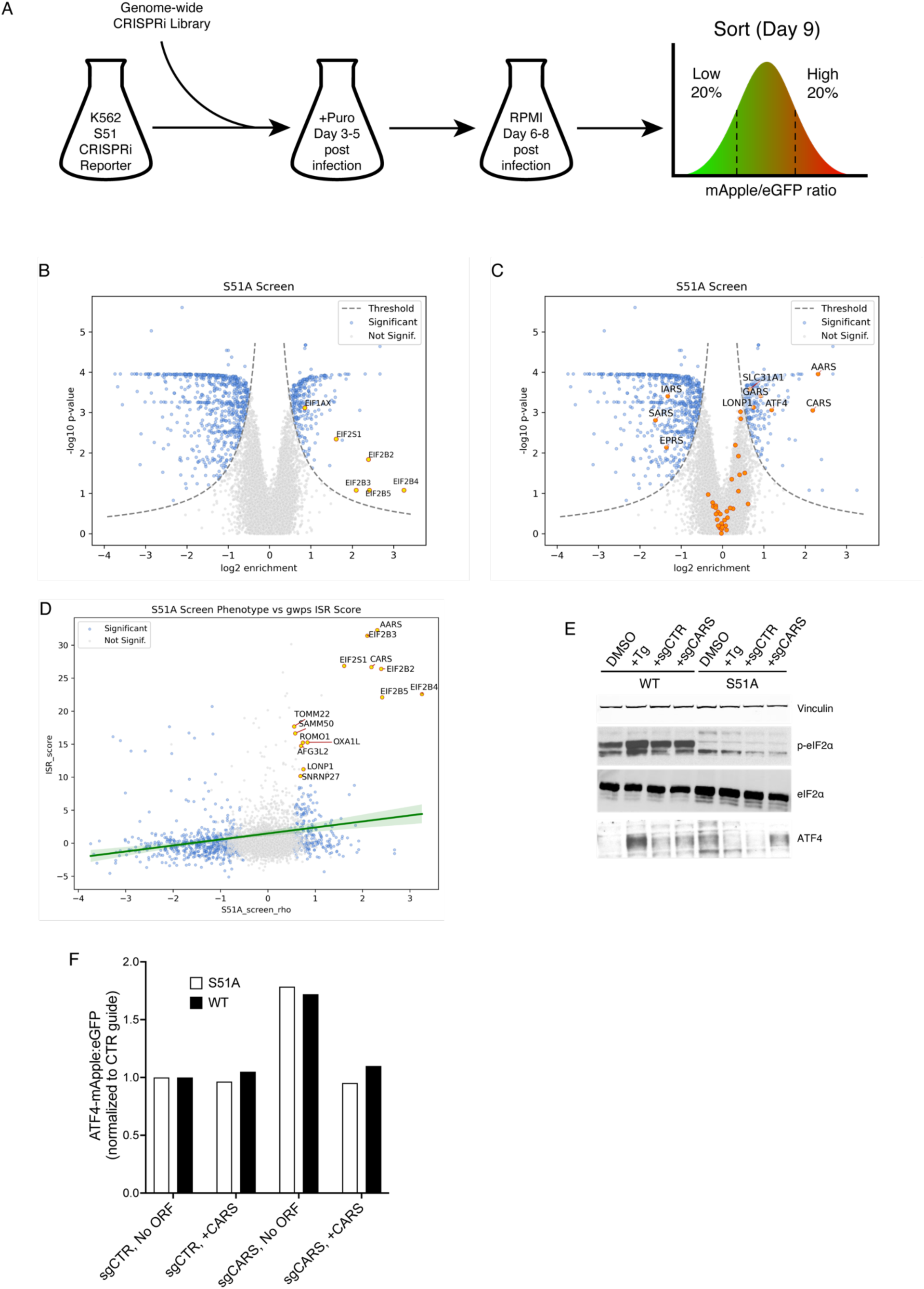
CRISPRi screen identifies CARS as an effector of ATF4 translation (A) Schematic of screen. (B) Screen results. Highlighted genes are all translation initiation machinery that enriched for reporter activity. (C) Screen results. Highlighted genes are top 30 genes for ISR activation in WT cells as defined in Methods. (D) Screen results compared to previously published ISR scores from Replogle et. al., 2022. (E) Immunoblot of ATF4, eIF2α, and p-eIF2α. Cells either expressed single sgRNAs, or were exposed to 75 nM thapsigargin or vehicle (DMSO) where indicated. (F) Expression of recombinant CARS is sufficient to suppress the ATF4 reporter induction caused by expression of a sgRNA targeting CARS.

When subsequently evaluating screen results, we focused on the enriched fraction of hit genes that were known to activate the ISR in wild type cells whose knockdown increased ATF4 reporter levels (Figure 2c). To do this, we re-analyzed data from genome-wide CRISPRi Perturb-seq studies to calculate ISR scores indicative of ATF4 activation in wildtype cells (Figure 2d), expecting that most ISR activating genes would do so in an eIF2α phosphorylation dependent manner, and therefore would not be enriched in our screen. To identify genes of interest, we then looked for genes with a high ISR score in the genome-wide Pertub-seq dataset that also activated the ATF4-mApple reporter in our screen.

The genes that scored highly in our screen for ATF4-mApple activation and also strongly activated the ISR in wild type cells fell into two categories. The first category was eIF2 machinery components as expected. The second category of strong hits that we chose to follow up on were the aminoacyl-tRNA synthetases cysteinyl-tRNA synthetase (CARS, gene name *CARS1*) and alanyl-tRNA synthetase (AARS, gene name *AARS1*) (Figure 2c,d). It has been previously established that loss of aminoacyl-tRNA synthetases strongly activates the ISR (Replogle *et al*., 2022b). This activation is hypothesized to occur in a GCN2-dependent manner as a result of depletion of cellular charged tRNA pools (Hinnebusch, 2005; Ishimura *et al*., 2016; Misra *et al*., 2021). However, we observed that while ATF4 reporter levels did not increase as a response to knockdown of the majority of aminoacyl-tRNA synthetases, *CARS1* and *AARS1* knockdown still resulted in reporter activity, suggesting some eIF2α phosphorylation independent pathway for at least these two tRNA-synthetases (Figure S2c). Additionally, it has been previously reported that cystine and cysteine transport are important downstream effectors of the ATF4 transcriptome and the ISR (Torrence *et al*., 2021), suggesting that CARS and cysteine biology might generally be important for translational regulation of ATF4.

We validated that *CARS1* knockdown increased translation of the ATF4 reporter and endogenous ATF4 without affecting phosphorylation of eIF2α via western blot (Figure 2e). This knockdown dependent ATF4 phenotype was suppressed by co-expression of recombinant CARS protein (Figure 2f, S2d), indicating that the increase in ATF4 translation was caused by loss of the native CARS protein. Similarly, individual knockdown of other selected genes recapitulated phenotypes observed in our genome-wide CRISPRi screen (Figure S2e, S2f).

### mTOR signaling is not required to drive ATF4 activation due to *CARS1* knockdown

Our screen results also indicated that mTOR-ATF4 signaling remained functional in our cell line (Figure 3a). They additionally suggested that activation of mTOR signaling via loss of TSC1 or TSC2, two suppressors of MTOR kinase activity (Figure 3a), led to increased ATF4 translation, a result in agreement with previously reported literature (Park *et al*., 2017; Replogle *et al*., 2022b). Western blotting for phosphorylated p70-S6 kinase, a known effector of mTOR signaling, indicated a modest increase in MTOR activity in both WT and S51A cells upon *CARS1* knockdown and in WT cells only upon thapsigargin treatment (Figure 3b). To examine if the *CARS1* knockdown phenotype was MTOR dependent, we pre-treated cells with the small molecule MTOR inhibitors rapamycin or torin-1 to inhibit mTOR signaling and then knocked down CARS. In both WT and S51A cells, *CARS1* knockdown still resulted in increased ATF4 translation, suggesting that this response, which we refer to henceforth as “CARS-ATF4”, is not dependent on mTOR signaling (Figure 3c).

**Figure 3:**
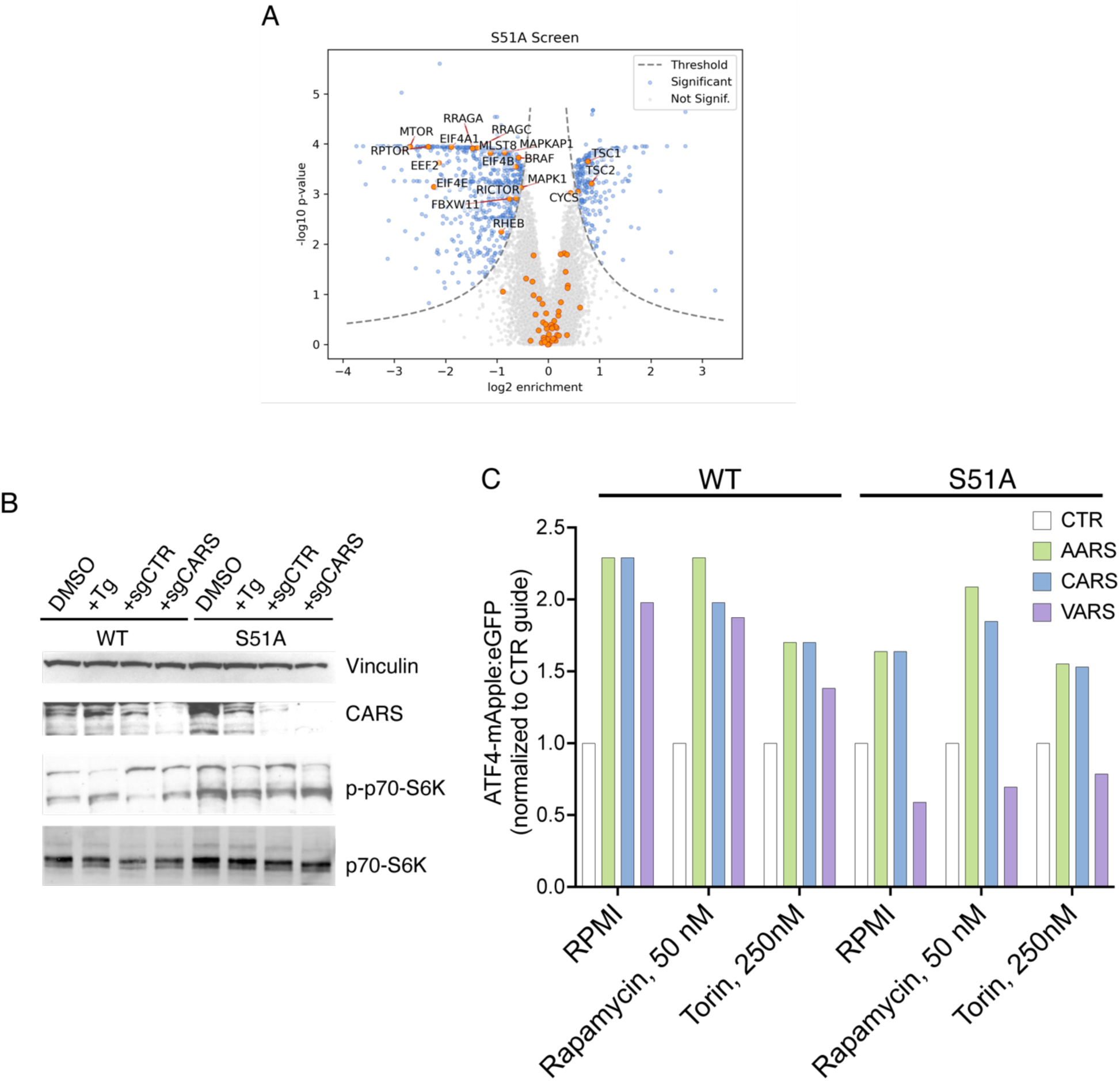
mTOR signaling is not required for ATF4 activation due to CARS1 knockdown (A) Screen results. Highlighted genes are genes involved in mTORC1 signaling as defined in Methods. (B) Immunoblot of CARS, p70-S6K, and p-p70-S6K. Cells either expressed single sgRNAs, or were exposed to 75 nM thapsigargin or vehicle (DMSO) where indicated. (C) Cells were cultured in rapamycin or torin 1 immediately post-infection with viral construct for expressing indicated sgRNA. Cells were maintained in drug or RPMI for 96 hours before measuring reporter levels by flow cytometry.

### CARS-ATF4 activation is potentially dependent on METAP2

To identify additional factors involved in ATF4 activation due to loss of CARS, we conducted a second genome-wide epistasis CRISPRi screen using a staggered infection strategy (Figure 4a, 4b). By first infecting cells with a genome-wide library and subsequently infecting with a CARS sgRNA, we aimed to identify genes required for CARS-ATF4 activation. In particular, we were interested in genes that either mildly activated or did not activate ATF4 in our initial screen of the S51A cell line, but suppressed *CARS1* knockdown mediated ATF4 activation.

**Figure 4:**
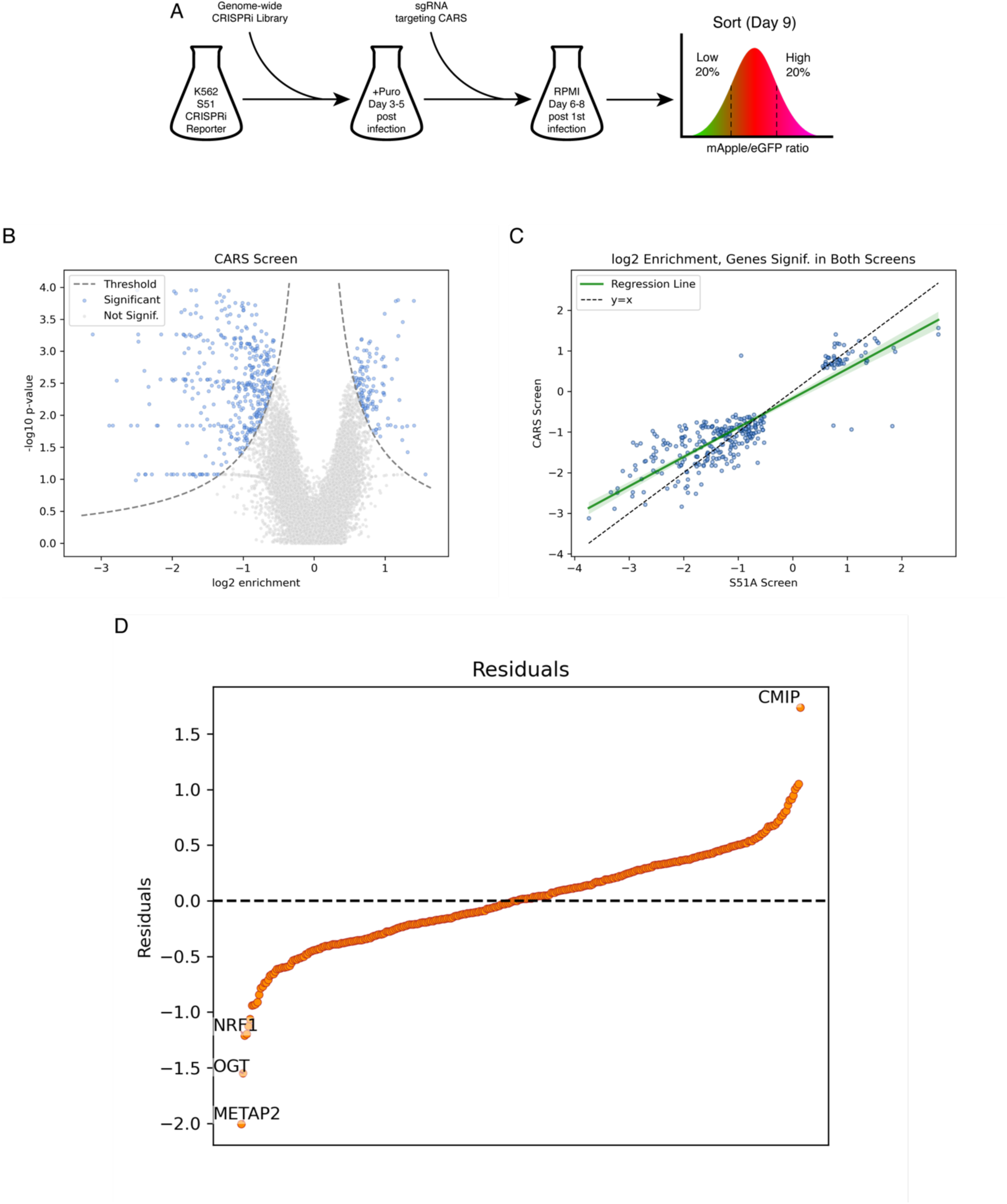
CARS-ATF4 activation is potentially dependent on METAP2 (A) Schematic of screen. (B) Screen results. (C) Comparison of phenotype scores between screen from Figure 2a and Figure 5a. Plotted points are all genes returning a statistically significant phenotype in at least one of the two screens, with linear regression model plotted. (D) Genes returning a statistically significant phenotype in at least one of the two screens, ordered by residual. Residuals are calculated using linear regression model from Figure 5c.

We reasoned that most genes in our two screens would have phenotype scores that correlated in a linear fashion. After running a linear regression model on phenotype scores for all genes that had significant phenotypes in at least one of our two screens (Figure 4c), we sorted all genes by residual CARS phenotype score. The top two genes after performing this analysis were methionine aminopeptidase 2 (METAP2) and *O*- linked *N*-acetylglucosaminyltransferase (OGT) (Figure 4d).

METAP2 is a metalloprotease responsible for the removal of the N-terminal methionine from newly synthesized proteins (Chiu *et al*., 2014). This process of N-terminal methionine excision is crucial for the maturation, stability, and functionality of many proteins (Sin *et al*., 1997; Griffith *et al*., 1998), and therefore METAP2 plays a key role in regulating protein turnover and translation rates (Tucker *et al*., 2008). METAP2 was originally identified as a 67-kilodalton protein that copurified with eIF2α (Datta *et al*., 1988; Ray *et al*., 1993), and this METAP2-eIF2α interaction was found to inhibit eIF2α phosphorylation in an *O*-linked β-*N*-acetylglucosamine (*O*-GlcNac) dependent manner (Datta *et al*., 1989; Gil *et al*., 2000).

OGT is the enzyme responsible for the post-translational addition of *O*-GlcNAc sugar groups onto serine and threonine residues of proteins (Haltiwanger *et al*., 1992; Lazarus *et al*., 2011). This *O*-GlcNAcylation is a reversible post-translational modification that regulates diverse cellular processes, including transcription, translation, signal transduction, and protein degradation (Lubas *et al*., 1997; Levine and Walker, 2016). OGT-mediated *O*-GlcNAcylation is regulated by the availability of UDP-GlcNAc, the substrate for OGT, and is therefore sensitive to various cellular stressors, signaling pathways, and bioavailability of precursor molecules (Hart *et al*., 2011).

Based on these facts, we believe that our screen results suggest a possible role for METAP2 in regulating the CARS-ATF4 response in an eIF2α phosphorylation independent manner that is dependent on the *O*-GlcNac modification state of METAP2.

## Discussion

Our findings build upon the body of literature establishing that ATF4 translation is not singularly regulated by eIF2α. Here, we establish that knockdown of the cysteinyl-tRNA synthetase *CARS1* is sufficient to activate ATF4 translation independent of eIF2α phosphorylation, and that this ATF4 translation is not a result of loss of charged cysteine-tRNA, but rather some yet to be determined non-canonical function of the CARS protein. Additionally, we show that this transcriptional response is dependent on the METAP2 and OGT genes, suggesting the existence of an until now unreported pathway for activating ATF4 translation.

Given lack of a known cytosolic sensor for cysteine levels in mammalian cells, it is possible that CARS, which depends on cysteine as a substrate, may play a role in sensing cysteine levels and directly regulating protein synthesis rates via modulation of ATF4. Given our screen results and the fact that METAP2 is known to directly interact with translation initiation machinery in an *O*-GlcNAc dependent manner, we believe that METAP2 suppresses ATF4 translation, and is required for signaling cysteine availability from CARS to translation machinery. A decrease in cysteine levels would then be sensed by CARS and result in inhibition of METAP2 and activation of ATF4 translation. We also believe that this interaction is dependent on OGT, possibly relying on CARS activated *O*-GlcNacylation of METAP2 by OGT to maintain METAP2 binding to translation initiation machinery and repression of ATF4 translation (Figure 5).

**Figure 5:**
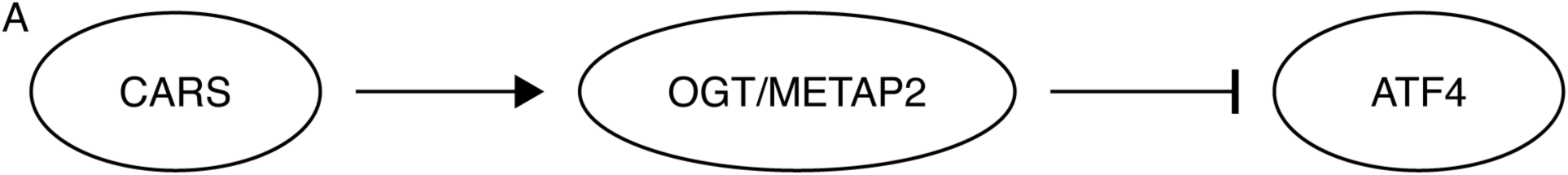
A proposed model of the CARS-METAP2/OGT-ATF4 pathway (A) CARS dependent activation of METAP2 is required to maintain METAP2 dependent repression of ATF4. We are unable to order the relationship between OGT and METAP2 based on our data but hypothesize that OGT acts upstream of METAP2 to regulate the *O*-GlcNAcylation state of METAP2. We also hypothesize but cannot support with data that METAP2 dependent repression of ATF4 is mediated by interactions between METAP2 and translation initiation machinery.

Our results also suggest that AARS may play a similar role in sensing cellular alanine levels, although we can only speculate as to whether AARS dependent signaling also requires METAP2 or OGT.

In general, our screen results suggest that there are more pathways capable of regulating ATF4 translation than previously realized. While our screen results suggest the possibility of immediately studying a large number of genes, the construction of the reporter system used in this study means that any follow up work will require initial arrayed validation of any hits to ensure that phenotypes observed in the screen are a result of changes in ATF4-mApple levels and not eGFP. Furthermore, given the fact that S51A cells only show a modest level of basal ATF4-mApple expression, coupled with the much lower magnitude of negative ISR scores in genome-wide Perturb-seq datasets compared to positive ISR scores, it is unclear if the strong phenotype scores returned by our screen correspond to real pathways of ATF4-mApple expression, or if the strong phenotypes are an artifact. We suggest that further examination of these screen results begins with study of hits that increased ATF4-mApple expression, and that any subsequent studies keep these caveats in mind.

An additional biological pathway observed in our initial screen the S51A cell line was the cholesterol biosynthesis pathway. Three strong hit genes *FDFT1*, *FDPS*, and *IDI1* represented three consecutive steps in cholesterol biosynthesis, with *FDFT1* controlling the rate-limiting step (Rudney and Sexton, 1986; Platt *et al*., 2014; Cerqueira *et al*., 2016). Cholesterol as a nutrient input is known to be sensed by the mTOR signaling pathway (Castellano *et al*., 2017), but there is no known pathway for direct cholesterol biosynthesis dependent stress sensing, with lipid bilayer fluidity changes as a result of cholesterol levels known to be signaled to the ISR as general ER stress via PERK (Schekman, 2013; Goldstein and Brown, 2015). However, the cholesterol biosynthesis dependent activation of ATF4 translation only replicated in one clone of our S51A cell line and so we chose not to further pursue this pathway. Despite this, there may still be some cholesterol dependent pathway for activating ATF4 independent of eIF2α phosphorylation that until now has been hidden by the ISR.

Increasing clinical interest in disease-state and context dependent activation or inhibition of the ISR and ATF4 translation will require more detailed understanding of different stress conditions and how they are responded to. The CARS-METAP2/OGT-ATF4 pathway we describe here is a potential therapeutic target to activate ATF4 translation in cells. It will be important to further clarify the mechanism of how this pathway activates ATF4 translation, as well as how this pathway impinges on or overlaps with other modes of activating ATF4 translation. Follow-up work can also focus on defining the exact transcriptional response driven by this pathway, and how it compares to other previously defined ATF4-dependent transcriptional responses. Additional exploration of these molecular pathways could help achieve beneficial clinical outcomes via fine-grained control of cellular responses to different inputs.

## Materials and Methods

### Antibodies

For a list of primary antibodies used in this study, see table 1.

**Table 1:** Primary antibodies used in this study.

| Antibody | Source and Catalog Number |
| --- | --- |
| ATF4 (rabbit mAb) | Cell Signaling Tech., #35031 |
| CARS, CARS-HaloTag7 (rabbit polyclonal) | Abcam, #ab235536 |
| eIF2α (rabbit mAb) | Cell Signaling Tech., #5324 |
| phospho-eIF2α Ser51 (rabbit mAb) | Cell Signaling Tech., #3398 |
| GAPDH (mouse mAb) | Cell Signaling Tech., #97166 |
| p70-S6K (rabbit mAb) | Cell Signaling Tech., #34475 |
| phospho-p70-S6K Thr389 (rabbit mAb) | Cell Signaling Tech., # 9234 |
| Vinculin (rabbit mAb) | Cell Signaling Tech., #13901 |

### Cell culture and lentiviral production

K562 cells were grown in RPMI-1640 containing 25mM HEPES, 2.0 g/L NaHCO3, 0.3 g/L L-Glutamine supplemented with 10% FBS, 2 mM glutamine, 100 units/mL penicillin and 100 mg/mL streptomycin. Cells were maintained between 0.25 x 106 - 1 x 106/ml. HEK293T cells were grown in Dulbecco’s modified eagle medium (DMEM) containing 10% FBS, 2 mM glutamine, 100 units/mL penicillin and 100 mg/mL streptomycin. All cell lines were grown at 37°C and 5% CO2. Serum starvation was performed by spinning cells at 300g for 5 minutes, washing twice in PBS, and resuspending in DMEM containing 2 mM glutamine, 100 units/mL penicillin and 100 mg/mL streptomycin but no FBS.

For lentivirus production, HEK293T cells were co-transfected with transfer plasmids (pCMV-VSV-G and delta8.9, Addgene #8454) and standard packaging vectors using the TransIT-LT1 Transfection Reagent (Mirus, MIR 2306). Supernatant was collected 48 hours post transfection, centrifuged, aliquoted and flash frozen at −80C until further use. Virus for the genome-wide CRISPRi screens was also generated using this method with one exception – HEK293T cells were seeded in IMDM (Thermo Fisher Scientific #1244053) supplemented with 20% inactivated fetal bovine serum (GeminiBio #100-106). In all instances, virus was rapidly thawed prior to transfection.

### Cell line generation

Generation of K562 CRISPRi lines with the ZIM3 KRAB domain has been previously described (Replogle *et al*., 2022a). The K562 CRISPRi cell line containing the ATF4 reporter (cAX10) was produced via co-infection of plasmids pXG237 and pXG260 (Guo *et al*., 2020). 5 Days post infection, cells were then treated with 75nM Thapsigargin and sorted for reporter induction using a Sony Cell Sorter (SH800S) to generate a polyclonal cell line. To generate a reporter cell line containing a stable homozygous S51A mutation in the endogenous *EIF2S1* locus (cAX23), cAX10 was nucleofected with a CRISPR-Cas9 RNP and a homology-directed repair template using the IDT Alt-R CRISPR-Cas9 System (http://www.idtdna.com/CRISPR-Cas9) targeting the endogenous *EIF2S1* locus. Nucleofection was done using a Lonza 4D-Nucleofector (AAF-1003X). Post nucleofection, cells were sorted as single-cell clones in 96-well plates using a Sony Cell Sorter (SH800S) and cells were tested for loss of reporter responsiveness to thapsigargin treatment. Homozygous mutations were confirmed after expansion by genomic PCR amplification with primers outside the endogenous 5’ and 3’ homology arms and sequencing (Sanger *et al*., 1977; Saiki *et al*., 1988).

See table 2 for a list of cell lines, RNA and DNA constructs, and primers used in this study.

**Table 2:**
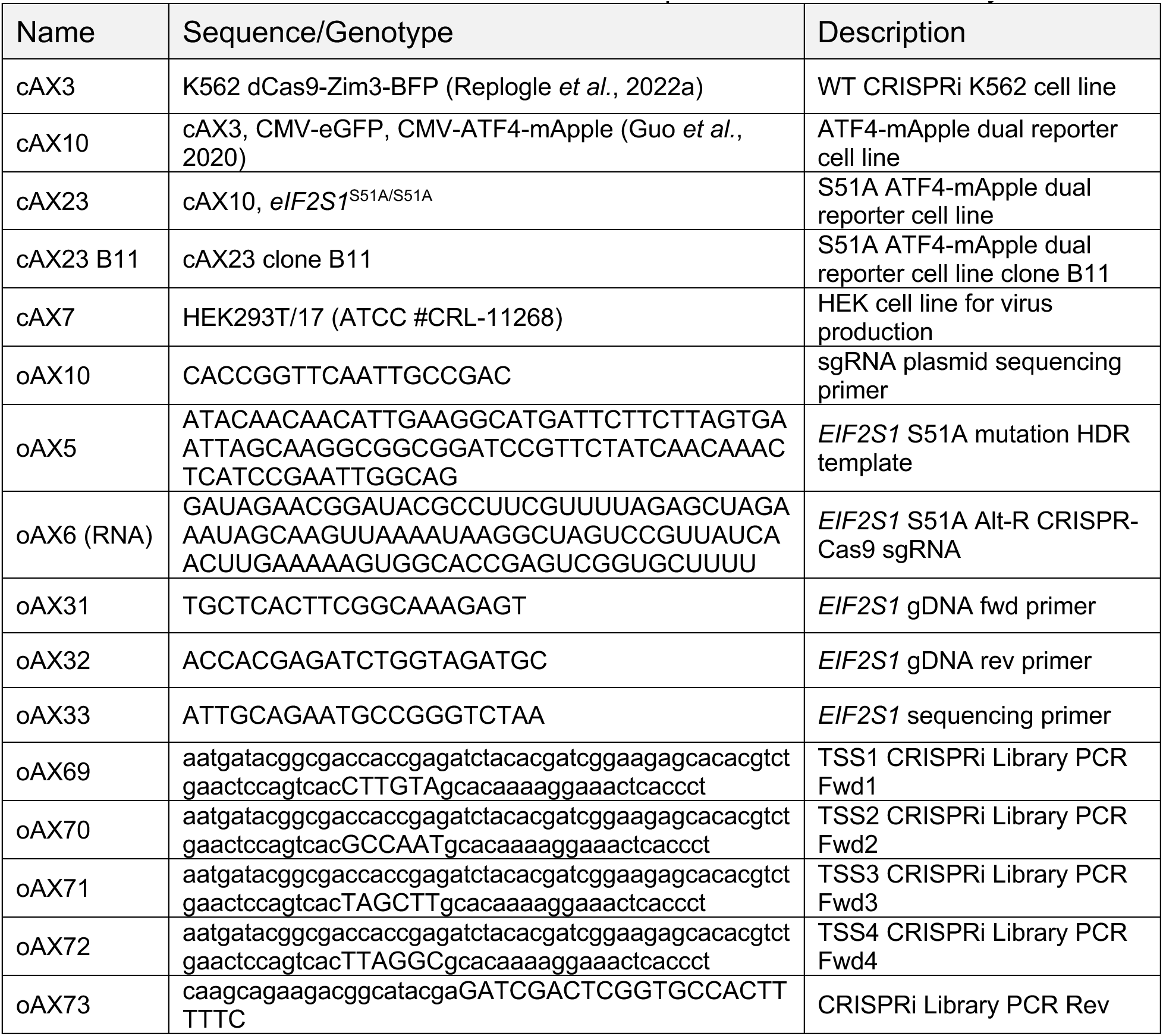
Cell lines, RNA and DNA constructs, and primers used in this study.

### Drug treatments

K562 cells and derived cell lines were diluted to 0.5 x 106/ml prior to drug treatment. An equal volume of RPMI with twice the final drug concentration was added. All durations of drug treatment are indicated in figures. To trigger ER stress, cells were treated with 75 nM of thapsigargin (Sigma-Aldrich #T9033), 20 μg/ml tunicamycin (Sigma-Aldrich # T7765), or 1 mM dithiothreitol (Thermo, #R0861). To inhibit mTOR activity, cells were treated with 50nM rapamycin (ThermoFisher #PHZ1235) or 250nM torin 1 (Tocris #4247).

### Constructs

Endogenous *CARS1* and *EIF2S1* sequences used in this study were sourced from UniProtKB/Swiss-Prot. All sequences were cloned into in vivo expression vectors using gene blocks synthesized by IDT (Integrated DNA Technologies, Coralville, IA). For cell rescue experiments, the CARS orf with a C-terminal HaloTag7 tag driven by the SFFV promoter was cloned into a lentiviral pU6-sgRNA, pEF-1α:Puro-T2A-GFP vector digested with NsiI/SbfI (Replogle *et al*., 2022b) by Gibson assembly.

To generate the construct for the second infection of the CARS screen, HaloTag7 was cloned into the lentiviral pU6:sgRNA, pEF-1α:Puro-T2A-GFP construct digested with NheI/EcoRI by Gibson assembly.

Single sgRNA constructs were generated by annealed oligo cloning of top and bottom oligonucleotides (Horlbeck *et al*., 2016) synthesized by IDT into the lentiviral pU6:sgRNA, pEF-1α:Puro-T2A-BFP or pU6:sgRNA, pEF-1α:HaloTag7 vector digested with BstXI/BlpI (Addgene, #84832). To deplete multiple genes at once we used a programmed dual sgRNA guide vector (Addgene #140096) (Replogle *et al*., 2022a).

See table 3 for list of single guides, dual guide combinations and respective protospacer sequences used in this study.

**Table 3:**
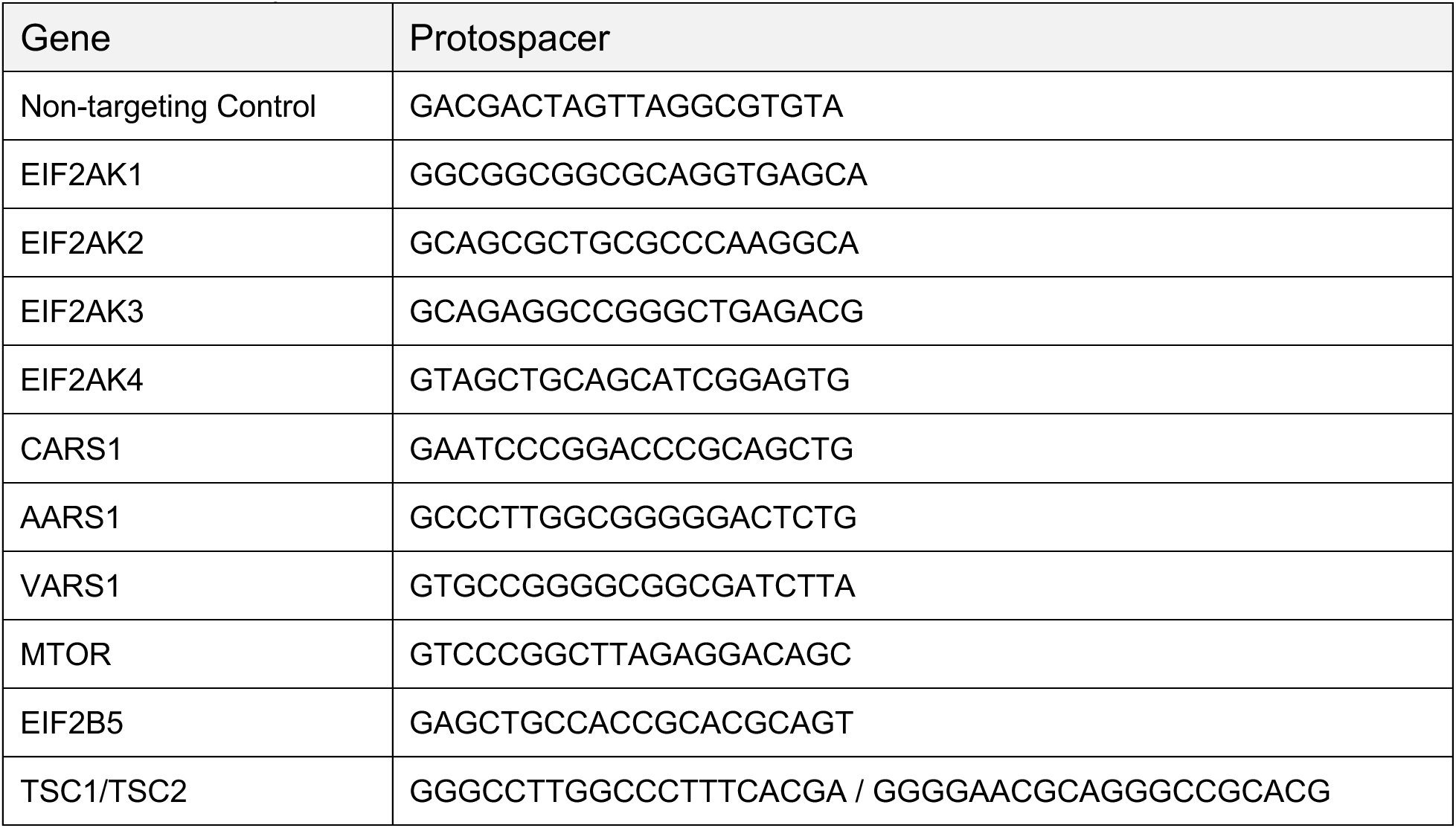
Single guides, dual guide combinations and respective protospacer sequences used in this study.

See table 4 for a list of plasmids and plasmid backbones used in this study.

**Table 4:** Plasmids used in this study.

| Plasmid | Description |
| --- | --- |
| pAX7 | Lentiviral pmU6:sgRNA, pEF-1α:Puro-T2A-BFP |
| pAX101 | Lentiviral phU6:sgRNA, pmU6:sgRNA, pEF-1α:Puro-T2A-BFP |
| pAX231 | Lentiviral pmU6:sgRNA, pSFFV:CARS-HaloTag7 |
| pTMB95 | Lentiviral pmu6:sgRNA, pEF-1α:HaloTag7.dna |
| pXG237 | Lentiviral pCMV:ATF4-mApple (Addgene #141281) (Guo *et al.*, 2020) |
| pXG260 | Lentiviral pCMV:EGFP (Addgene #141282) (Guo *et al.*, 2020) |

### CRISPRi screens

Genome-scale FACS-based CRISPRi screens were performed as previously described (Gilbert et. al, 2014, Horlbeck et. al, 2016). All screens were performed in duplicate. For each, the hCRISPRi-V2 compact library (Addgene #83969, 5 sgRNAs/TSS) was transduced at a MOI (multiplicity of infection) <1 into 400 million cells. The percentage of cells transduced was measured at 48 hours post transduction by recording the percent of BFP positive cells (∼20-30%). Replicates were grown separately in 1L of RPMI-1640 in 3L spinner flasks (Bellco, SKU: 1965-61030).

For the S51A screen, cells were selected with 1 ug/ml puromycin starting at 48 hours post-transduction for 3 days until the transduced cells accounted for 90-95% of the population, and then maintained at 0.5 x 10^6^/ml to ensure an average coverage of more than 1000 cells/sgRNA. For sorting on a BD FACS Aria II, cells were gated for BFP (indicating guide-positive cells) and on the RFP:GFP ratio (highest and lowest 20%). Approximately 40 million cells each were collected from the top and bottom 20% based on the RFP:GFP ratio, pelleted and flash frozen.

For the CARS screen, cells were selected with 1 ug/ml puromycin starting at 48 hours post-transduction for 3 days until the transduced cells accounted for 90-95% of the population, and then maintained at 1.0 x 10^6^/ml to ensure an average coverage of more than 2000 cells/sgRNA. 24 hours after the last puromycin treatment, cells were transduced with virus containing the sgRNA targeting CARS and HaloTag7 at a MOI (multiplicity of infection) <1. After transduction, cells were maintained at 0.5 x 10^6^/ml to ensure an average coverage of more than 1000 cells/sgRNA. For sorting on a BD FACS Aria II, cells were treated with HaloTag-JF646 (Grimm *et al*., 2015) and gated for BFP (indicating guide-positive cells), JF646 (indicating sgCARS positive cells), and on the RFP:GFP ratio (highest and lowest 20%). Approximately 20 million cells each were collected from the top and bottom 20% based on the RFP:GFP ratio, pelleted and flash frozen.

Genomic DNA was isolated from frozen cells using the Nucleospin Blood XL kit (Takara Bio, #740950.10) and amplified with barcoded primers by index PCR (see for sequences). Guide libraries (∼264 bp) were purified using SPRIbeads (SPRIselect Beckman Coulter #B23318) followed by sequencing with an Illumina HiSeq2500 high throughput sequencer.

### Processing of CRISPRi screen data

Using the Python-based ScreenProcessing pipeline, sequencing reads were aligned to the CRISPRiv2 library sequences, counted and quantified. Negative control gene generation and calculation of phenotypes and Mann-Whitney p values was performed as previously described (Gilbert *et al*., 2014; Horlbeck *et al*., 2016). Phenotypes from sgRNAs targeting the same gene were collapsed into a single mApple:GFP ratio phenotype derived from the average of the top three scoring sgRNAs (by absolute value) and assigned a p-value through the Mann-Whitney test of all sgRNAs targeting the same gene compared to the non-targeting controls. All additional CRISPRi screen data analyses were performed in Python 3.9.12 using a combination of Numpy (v1.19.0), Pandas (v0.25.0), and Scipy (v1.5.4). Volcano plots were generated using a combination of the Python Bokeh package (v3.1.1) and Seaborn (v0.13.2). Gene lists of top activators of pathways were taken from genome-wide PERTURBseq studies (Replogle *et al*., 2022b).

### Flow cytometry

For all reporter experiments, sgRNA constructs were individually packaged into lentivirus, spinfected into K562 cells and analyzed by flow cytometry. For experiments testing the effects of single or dual sgRNAs on reporters, sgRNA packaged into lentivirus was first spinfected into K562 cells at a MOI <1 (∼20-40%). Transduced cells were selected with puromycin at 1 ug/ml for 3 days starting at 48 hours post-spinfection, Where indicated, cells were sorted by % BFP-positive concurrently with flow cytometry analysis to select for cells either expressing guides for downstream qRT-PCR, immunoblotting, RNAseq, and tRNAseq analyses. All flow cytometry data was collected on an NXT Flow Cytometer (Thermo Fisher). Data analysis was performed in Python using the FlowCytometryTools package and a combination of Pandas, Numpy, Scipy and Seaborn libraries, or FlowJo (v10.10). Reporter levels induction levels were measured as the mean mApple/eGFP ratio normalized to an experimental control sample.

### SDS-PAGE and western blotting

Cells (approx. 0.5 - 1×10^6^ per sample) were first washed with PBS and pelleted, and then lysed in RIPA buffer (Sigma-Aldrich, R0278) and 1x Pierce complete protease and phosphatase inhibitor cocktail (Thermo Fisher, A32959). Lysates were clarified to remove cell debris by rotating end-over-end at 4°C briefly and subsequently centrifugated at 5,000g for 5 minutes at 4°C. Total protein material for each sample was quantified using the Pierce BCA Protein Assay kit (Thermo Fisher, 23225) for equal loading (Lowry *et al*., 1951; Bradford, 1976; Smith *et al*., 1985). Samples were first boiled with 6x Laemmli SDS buffer (Laemmli, 1970) at 95°C for 5 minutes and then proteins were separated on Bolt 4-12% Bis-Tris gels (Thermo Fisher, NWO4122BOX and NWO4127BOX), transferred to Nitrocellulose membranes using the Mini Trans-Blot Cell (BioRad) kit at Mixed MW (molecular weight) settings according to the manufacturer’s instructions (Towbin *et al*., 1979), blocked with EveryBlot Blocking Buffer (BioRad #12010020) at room temperature for 1 hour, and subsequently probed with relevant primary antibody. Secondary antibodies used were goat-anti-mouse-HRP and goat-anti-rabbit-HRP (Biorad, #172-1011 and #170-6515).

### Statistics

Significance values (where applicable) were set to 0.05. Experiment specific details can be found in respective figure legends. All error bars indicate mean ± s.d., n = 3.

## Acknowledgements

We thank M. Kampmann for sharing the ATF4 screening system using the dual reporter strategy. We thank Y.H. Chen, G. Muthukumar, and K. E. Yost for assistance with computational analyses. We thank T.M. Bertozzi for sharing the pU6:sgRNA, pEF-1α:HaloTag7 construct. We thank G. Muthukumar, Z. Levine, and all other members of the Weissman lab for their discussion regarding this work. We thank the Whitehead Institute and Koch Institute Flow Cytometry Cores for access to FACS machines, and the Whitehead Institute Genome Technology Core for support with sequencing of screen libraries. This work was supported by the Howard Hughes Medical Institute (J.S.W.).

## Author Contributions

A.E.X. and J.S.W. were responsible for the conception, design and interpretation of experiments and wrote the manuscript with input from all authors. A.E.X. led and performed all experiments and accompanying data analysis. J.S.W. funded the study. J.S.W. is an investigator of the Howard Hughes Medical Institute.

---

**Supplemental Figure 1:**
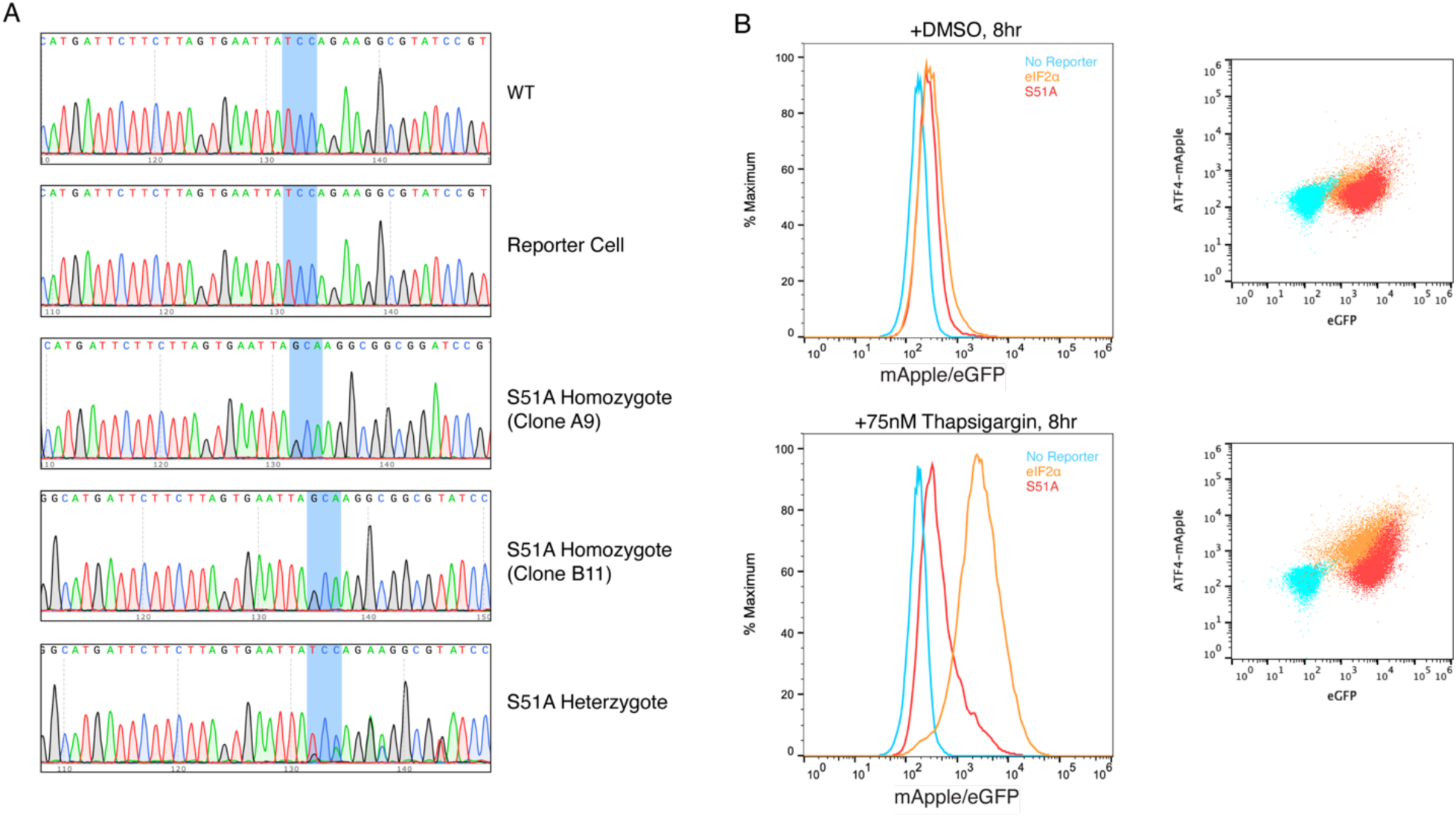
Generation of homozygous S51A alleles, related to Figure 1 (A) Sanger sequencing traces of different cell lines. Highlighted codon indicates the serine 51 residue, with heterozygous alleles showing as peaks with signal for multiple nucleotides. (B) Flow cytometry data corresponding to Figure 1B.

**Supplemental Figure 2:**
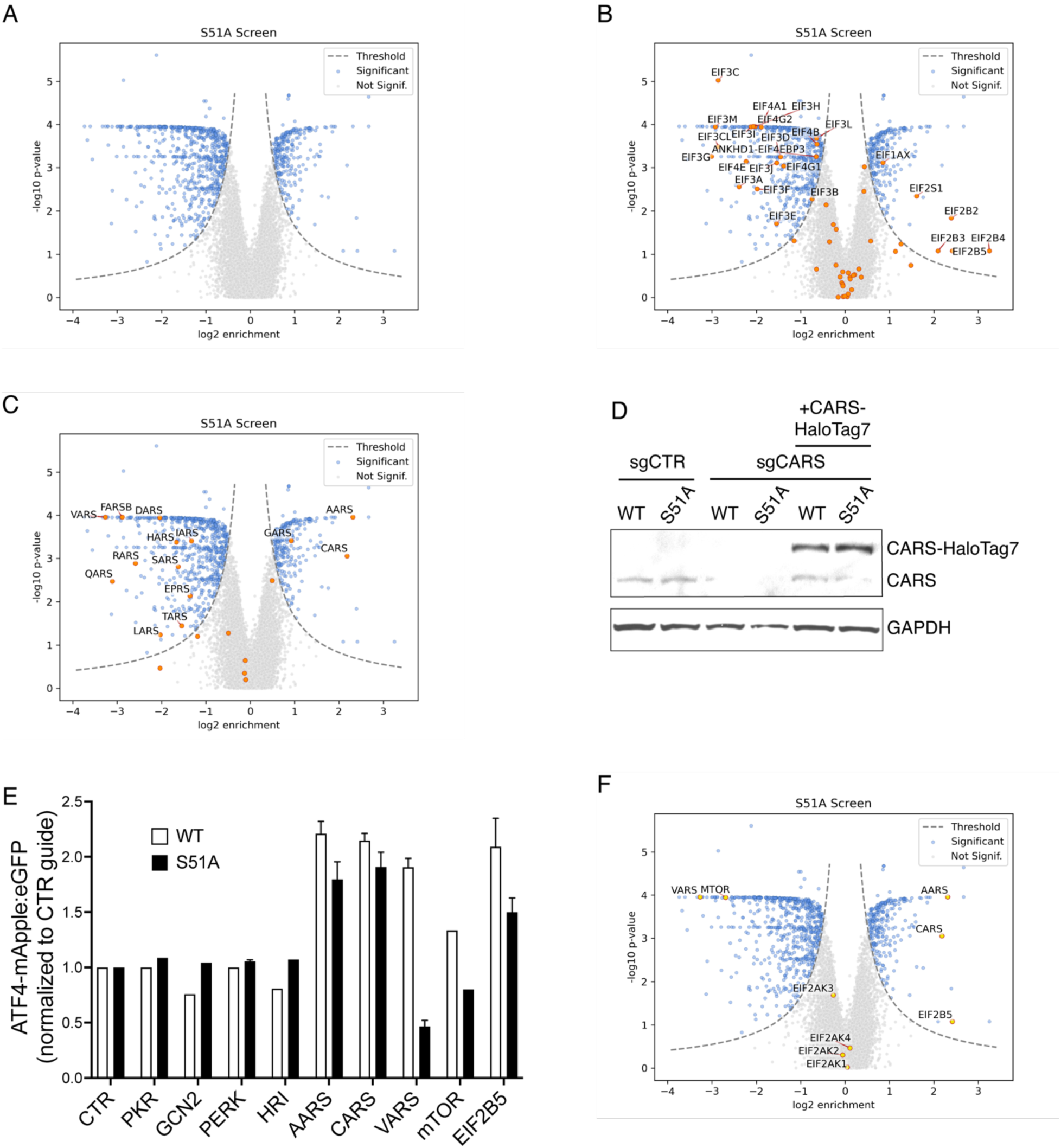
Additional S51A CRISPRi screen validation, related to Figure 2 (A) Screen results for screen in Figure 2a. (B) Screen results. Highlighted genes are all translation initiation machinery. (C) Screen results. Highlighted genes are all aminoacyl-tRNA synthetases. (D) Immunoblot of cells expressing single RNAs for CARS and recombinant CARS-HaloTag7. (E) WT and S51A reporter cells expressing single sgRNAs before measuring reporter levels by flow cytometry. (F) Screen results. Highlighted genes are genes from Supplemental Figure 2e.

